# Sequential Dual-Ion MALDI Glycotyping Enables Rapid Phenotypic O-Antigen Typing of *Escherichia coli* and *Shigella*

**DOI:** 10.64898/2026.03.20.713141

**Authors:** Shogo Urakami, Hiroshi Hinou

## Abstract

Accurate O-antigen typing of Gram-negative bacteria is important for surveillance, outbreak investigation, and quality control of reference strains. However, commonly used typing approaches, including serological agglutination assays and molecular methods, do not always resolve structural variation in expressed O-antigen phenotypes. Here, we describe an improved MALDI glycotyping workflow based on matrix-assisted laser desorption/ionization time-of-flight mass spectrometry (MALDI-TOF MS) that enables rapid phenotypic characterization of O-antigen repeating units (RUs). The workflow uses a sequential dual-ion acquisition strategy in which positive-ion spectra are obtained first and negative-ion analysis is triggered only when RU signals are absent, enabling detection of both neutral and acidic O-antigen structures from the same sample spot. Applied to a diverse panel of 71 *Escherichia coli* and *Shigella* strains, RU-derived signals were detected in more than 80% of isolates. The approach resolved modification-level structural variation and discriminated isobaric O-antigen phenotypes, enabling scalable phenotypic profiling of O-antigen composition and inference of candidate O-antigen identities from RU-level information. Integration with agglutination testing further revealed discrepancies between archived serotype annotations and expressed O-antigen phenotypes, enabling reassignment of several strains to alternative O-antigen types. Because the workflow can be implemented on MALDI-TOF MS platforms already widely used for microbial identification, this method provides a practical phenotypic complement to conventional O-antigen typing in clinical microbiology laboratories and remains compatible with rapid single-colony MALDI workflows used in routine microbial identification.

**Importance:** Accurate O-antigen characterization is essential for pathogen surveillance and quality control of reference strain integrity. However, existing methods often rely on genetic or serological proxies rather than direct structural analysis of the expressed O-antigen. We established a sequential dual-ion MALDI glycotyping method that enables rapid phenotypic characterization of both neutral and acidic O-antigen repeating units in *E. coli* and *Shigella*. This approach detected O-antigen signals in over 80% of a diverse strain panel and identified fine structural modifications that conventional tests miss. Importantly, our method uncovered discrepancies in archived serotype data, allowing for the corrected reassignment of several reference strains. By integrating glycan phenotyping into existing MALDI-TOF MS workflows already common in clinical settings, this method offers a robust and scalable tool for high-resolution bacterial typing and quality control.

## Introduction

Bacterial O-antigens are structurally diverse polysaccharides that form the outermost domain of lipopolysaccharide in Gram-negative bacteria. In *Escherichia coli* and *Shigella*, O-antigen diversity provides the basis for serotype designation and remains important in clinical microbiology for pathogen surveillance, outbreak investigation, and quality assurance of reference strain collections (1-3). More than 180 O-antigen types have been described in *E. coli*, reflecting extensive variation in repeating unit (RU) sugar composition, linkage patterns, stereochemistry, and chemical modifications such as O-acetylation (4). Despite this importance, routine methods do not always provide sufficient phenotypic resolution for accurate characterization of expressed O-antigen diversity in clinical and epidemiological settings (1, 5).

Current O-antigen typing strategies rely mainly on serological agglutination assays and molecular approaches such as PCR-based analysis of O-antigen biosynthetic loci (6-8). Although these methods have enabled large-scale O-antigen typing, they are limited to predefined targets and may not fully resolve structural diversity beyond established serotypes. In particular, they can struggle to discriminate intra-serotype variants and modification-level differences and cannot readily capture previously uncharacterized phenotypes. These limitations highlight the need for complementary approaches that directly characterize expressed O-antigen phenotypes.

Matrix-assisted laser desorption/ionization time-of-flight mass spectrometry (MALDI-TOF MS) is already widely used in clinical microbiology laboratories for rapid species identification because of its speed, simplicity, and low cost (9-11). However, its routine use remains largely confined to protein mass fingerprinting. We recently introduced MALDI glycotyping, a strategy that enables direct detection of O-antigen repeating units (RUs) from a single bacterial colony within one hour, thereby expanding routine MALDI-TOF MS workflows beyond protein fingerprinting to enable glycan phenotyping (12, 13). Although this initial approach showed that RU composition could be directly measured to discriminate subtypes and previously unreported glycotypes, it was evaluated on a limited number of strains and primarily detected O-antigens composed of neutral RUs, leaving acidic O-antigen structures incompletely represented.

In this study, we establish an improved MALDI glycotyping workflow for routine phenotypic characterization of both neutral and acidic O-antigen structures in *Escherichia coli* and *Shigella*. The method uses sequential dual-ion mode acquisition, in which positive-ion spectra are obtained first and negative-ion measurements are triggered only when RU-derived signals are not detected. By integrating controlled chemical pretreatment with this conditional acquisition strategy, the workflow enables direct detection of structurally diverse O-antigen phenotypes from the same sample spot without additional sample handling or instrument modification. Using a diverse panel of 71 *E. coli* and *Shigella* strains representing multiple O-antigen serotypes, we evaluated the ability of this workflow to detect O-antigen repeating unit signals, resolve structural variation among related O-antigens, and identify discrepancies between archived serotype assignments and expressed O-antigen phenotypes. RU-derived signals were detected in more than 80% of strains, demonstrating the practical feasibility of MALDI glycotyping for scalable phenotypic characterization of O-antigen diversity in routine microbiology workflows.

## Materials and Methods

### Materials

Media for revival of L-dried specimens DAIGO and hipolypeptone were purchased from Nihon Pharmaceutical Co., Ltd. (Osaka, Japan). 2,5-Dihydroxybenzoic acid (DHB), sodium bicarbonate (NaHCO₃), sodium chloride (NaCl), acetonitrile (HPLC grade), magnesium sulfate heptahydrate, agar powder, potassium carbonate (K₂CO₃), and hydrochloric acid (HCl) were purchased from Fujifilm Wako Pure Chemical Industries, Ltd. (Osaka, Japan). Trifluoroacetic acid (TFA) was purchased from Watanabe Chemical Industry Co., Ltd. (Hiroshima, Japan). 1,5-Diaminonaphthalene (DAN) was purchased from Sigma-Aldrich Corp. (St. Louis, MO, USA). 3,4-Diaminobenzophenone (DABP) was purchased from Tokyo Chemical Industry Co., Ltd. (Tokyo, Japan). Bacto-yeast extract was purchased from BD Biosciences (San Diego, CA, USA). CHROMagar *E. coli* (CHROMagar, Paris, France) and Accudia SS agar (Shimadzu Diagnostics Corp., Kyoto, Japan) were used as selective media for bacterial isolation. All water used in this study was purified using a Milli-Q water purification system (Direct-Q 3 UV; Merck Millipore, Tokyo, Japan).

### Bacterial strains and culture conditions

*Escherichia coli* and *Shigella* strains used in this study are listed in Table S1. All strains were obtained from the Gifu Type Culture Collection of the Microbial Genetic Resource Stock Center, Gifu University Graduate School of Medicine (Gifu, Japan), via the National BioResource Project (Japan). Lyophilized strains were revived on agar plates prepared with Media for revival of L-dried specimens DAIGO. Prior to MALDI analysis, *E. coli* strains were streaked on CHROMagar *E. coli* and *Shigella* strains on Accudia SS agar for isolation and confirmation. Single colonies were then transferred to the nonselective DAIGO agar medium and cultured at 37 °C overnight before analysis.

### Sample preparation for MALDI glycotyping

Sample preparation for MALDI glycotyping was performed as previously described (12). Briefly, multiple bacterial colonies were suspended in 1 mL of water, washed twice by centrifugation at 15,000 × g for 2 min, and resuspended in 30 μL of water. The optical density was adjusted to 1.7–2.0. An aliquot (1.5 μL) was mixed with 0.5 μL of 400 mM HCl (final concentration, 100 mM), incubated at 90°C for 10 min, and centrifuged at 20,000 × g for 5 min. The supernatant (0.35 μL) was deposited onto a MALDI target plate, air-dried, overlaid with 0.35 μL of matrix solution, and dried at room temperature.

### MALDI matrix preparation

MALDI matrix solutions were prepared as follows. Stock solutions of DAN (50 mM) and DABP (50 mM) were prepared in acetonitrile/water (1:1, v/v). A stock solution of DHB (500 mM) was prepared in acetonitrile/water (9:1, v/v). NaHCO_3_ (100 mM) and K_2_CO_3_ (50 mM) were prepared in water.

Four matrix formulations were prepared: DAN/DHB/Na (14), DAN/DHB/K (13), DHB (15), and DABP (16). For the DAN/DHB/Na matrix, 2 μL of 500 mM DHB solution, 4 μL of 50 mM DAN solution, and 1 μL of 100 mM NaHCO_3_ were mixed and diluted to a final volume of 100 μL with acetonitrile/water (1:1, v/v). For the DAN/DHB/K matrix, NaHCO_3_ was replaced with 50 mM K_2_CO_3_ using the same volumes and dilution procedure. DAN/DHB/Na and DAN/DHB/K matrices were used within 12 h of preparation.

For the DHB matrix, 2 μL of 500 mM DHB solution was diluted to 100 μL with acetonitrile/water/trifluoroacetic acid (50:50:0.1, v/v/v). For the DABP matrix, 20 μL of 50 mM DABP solution was diluted to 100 μL using the same solvent system. All solutions were prepared using Milli-Q water.

All MALDI glycotyping analyses were performed using DAN/DHB/Na and DAN/DHB/K matrices. Unless otherwise indicated, comparative analyses presented in the main text and supplementary materials were based on spectra acquired with the DAN/DHB/K matrix. MS/MS analyses were performed using the DAN/DHB/Na matrix because it produced more abundant fragment ions and facilitated spectral interpretation.

### MALDI-TOF MS and MS/MS acquisition

Mass spectra were acquired using an Ultraflex III MALDI-TOF/TOF instrument (Bruker, Bremen, Germany) equipped with a 200-Hz Smartbeam Nd:YAG laser (355 nm). Spectra were acquired in reflectron mode with ions accelerated to 25.0 kV. The low-mass-ion deflector was set to 700 Da, and 1,000 laser shots were accumulated for each spectrum. MS/MS analyses were performed in LIFT-TOF/TOF mode with precursor ions accelerated to 8 kV in the ion source and 19 kV in the LIFT cell. Fragment ions were analyzed as metastable postsource decay (PSD) products without collision-induced dissociation.

### Sequential dual-ion acquisition workflow

For each sample spot, spectra were first acquired in positive-ion mode. Negative-ion mode acquisition was performed only when positive-ion mode spectra lacked RU-derived signals. Positive- and negative-ion mode spectra were acquired sequentially from the same sample spot without additional sample preparation or matrix exchange.

### RU signal assignment and concordance assessment

RU-derived signals were defined as ion series consistent with integer multiples of a single RU mass and reproducibly detected above baseline. For each strain, RU exact mass was inferred from the dominant RU-derived ion series, taking the observed adduct species into account. Monosaccharide composition and modification state were assigned primarily from positive-ion mode spectra, with MS/MS used selectively for confirmation. For strains with reported O-antigen assignments, concordance was assessed by comparison of MALDI-derived RU features with corresponding reference structures (4, 8, 17). Strains were classified as concordant, discordant, or not detected according to agreement between MALDI-derived RU exact mass, inferred monosaccharide composition, and observable modification state and the corresponding reference O-antigen structure.

### Structure-informed comparison with reported O-antigen structures

For strains with discordant or unresolved O-antigen assignments, MALDI-derived RU exact masses, inferred monosaccharide compositions, and observable modification states were compared with reported *E. coli* O-antigen structures. Reported K-antigen structures (18) were also reviewed to exclude capsular polysaccharides as sources of RU-derived signals. Candidate O-antigen identities were first screened by agreement of RU exact mass and then further narrowed using compositional and modification-level consistency. Comparative analysis was used to constrain compatible O-antigen candidates rather than to force definitive serotype assignment when structural ambiguity remained.

### Agglutination serotyping

Agglutination serotyping was performed using a commercially available *E. coli* O-antigen antisera panel (Denka Seiken Co., Ltd., Tokyo, Japan) according to the manufacturer’s instructions. Bacterial colonies were suspended in saline, heat-treated at 121°C for 15 min, centrifuged at 900 × g for 20 min, and resuspended in saline to an optical density of 1.7–2.0. Approximately 30 μL of antiserum and 10 μL of bacterial suspension were mixed on a glass slide and agitated for at least 1 min. Reactions were visually assessed and documented by microscopy.

## Results

### Sequential dual-ion MALDI glycotyping enables detection of neutral and acidic O-antigen repeating units

To evaluate the MALDI glycotyping workflow, we applied a sequential dual-ion mode acquisition strategy in which spectra were first acquired in positive-ion mode and negative-ion measurements were performed only when RU-derived signals were absent (Fig. 1). This strategy was illustrated using an *Escherichia coli* O53 (19) strain (GTC 03596), whose repeating unit (RU) contains a uronic acid residue and is therefore expected to be poorly detected in positive-ion mode. As expected, no RU-derived signals were observed under positive-ion conditions, whereas subsequent negative-ion mode acquisition yielded a clear [1RU − H]⁻ ion at *m/z* 859.4 (<u>Fig. S1.20; complete spectral atlas in Fig. S1</u>). These results demonstrate that sequential dual-ion mode acquisition enables selective detection of acidic O-antigen phenotypes that are inaccessible under fixed acquisition conditions.

**Figure 1.**
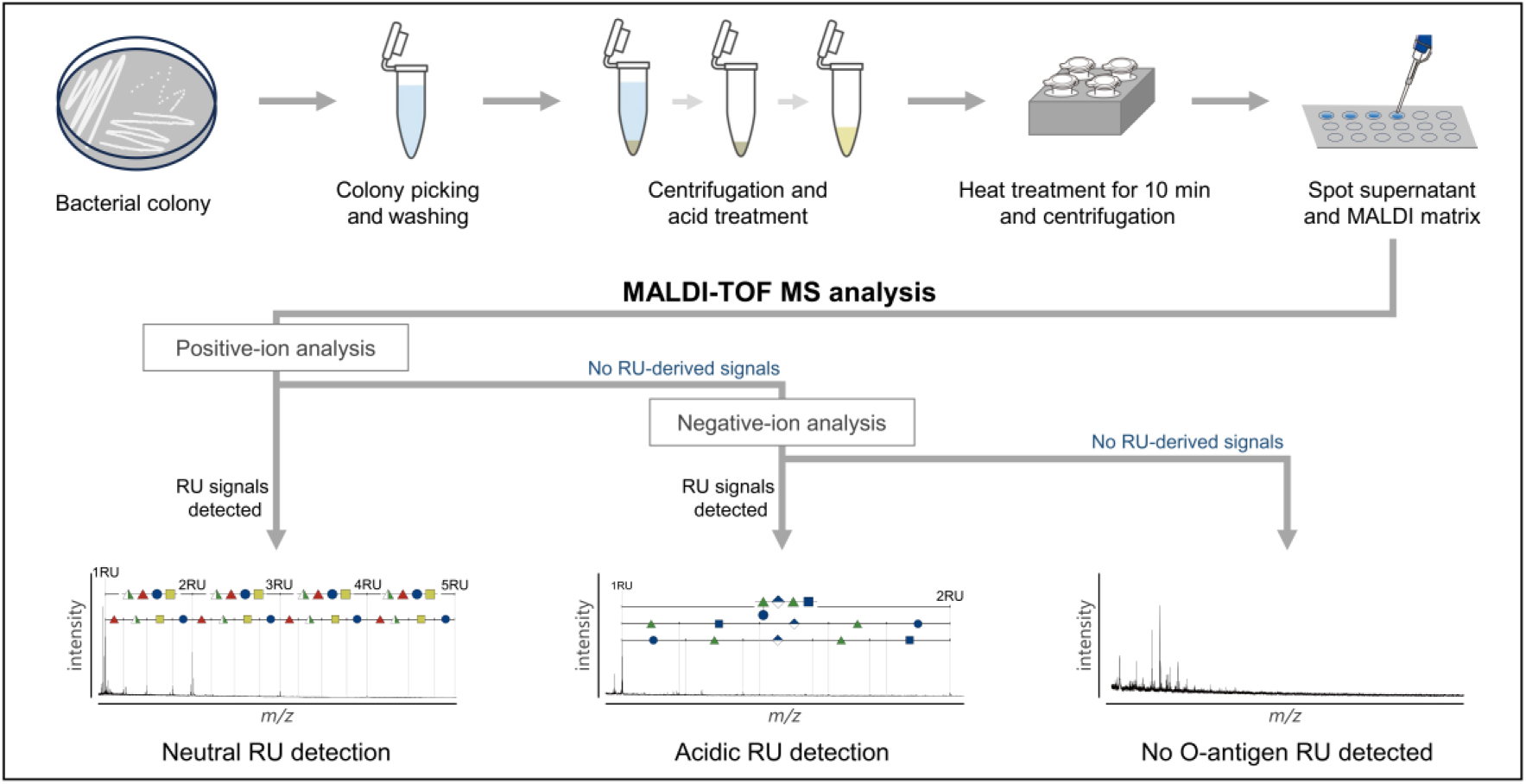
Sequential dual-ion MALDI glycotyping workflow for detection of neutral and acidic O-antigen repeating units. Bacterial colonies are processed by mild acid pretreatment followed by MALDI-TOF MS analysis to enable direct phenotyping of O-antigen repeating units (RUs). Samples are first analyzed in positive-ion mode to detect neutral RUs. When no RU-derived signals are observed, the same sample spot is subsequently analyzed in negative-ion mode. This conditional sequential dual-ion acquisition strategy enables detection of acidic O-antigen phenotypes, including species containing negatively charged residues such as uronic acids, that are often missed under fixed single-ion acquisition. Representative outcomes illustrate neutral RU detection, acidic RU detection, and no RU detection. Raw spectra for all strains are provided in <u>Fig. S1.1–S1.71</u>.

To evaluate matrix performance in negative-ion mode, we compared four matrix systems, including conventional glycan matrices (DHB and DABP) and two matrices previously developed for positive-ion glycotyping (DAN/DHB/Na and DAN/DHB/K). RU-derived signals were detectable with all matrices tested; however, DAN/DHB/Na and DAN/DHB/K provided substantially higher signal intensity and signal-to-noise ratios than DHB or DABP, enabling confident assignment of the O53 RU composition and RU mass (<u>Fig. S2</u>).

These results demonstrate that sequential dual-ion mode acquisition combined with optimized matrix systems is an effective strategy for MALDI-based phenotyping of both neutral and acidic O-antigen repeating units.

### Species-scale validation of MALDI glycotyping across *Escherichia coli* and *Shigella* strains

To assess the scalability of conditional MALDI glycotyping for O-antigen phenotyping, we applied the workflow across a diverse panel of 71 *Escherichia coli* and *Shigella* strains <u>(Fig. S1.1–S1.71 and Table S1</u>). These strains included multiple clinically relevant serotypes such as O157 and O26. Each strain was first analyzed in positive-ion mode, with negative-ion acquisition triggered only when positive-ion spectra lacked RU-derived signals (Fig. 1).

Using this conditional workflow, RU-derived signals were detected in 57 of the 71 strains (80.3%) (Fig. 2A). Detection patterns differed between species: most *E. coli* strains yielded RU-derived signals in positive-ion mode, consistent with the prevalence of neutral O-antigen compositions, whereas Shigella strains more frequently required negative-ion acquisition, reflecting the higher occurrence of acidic O-antigen components. Across all strains, individual isolates typically yielded multiple RU-derived ion series in positive-ion mode and one or two series in negative-ion mode (Fig. 2B and C). Detected RUs ranged from di- to hexasaccharides, with pentasaccharide units being most prevalent (Fig. 2D). These RU-derived signals collectively reflected substantial compositional diversity, encompassing twelve distinct monosaccharide residues and seven non-sugar constituents together with variable RU sizes and modification states (Fig. 2D and E).

**Figure 2.**
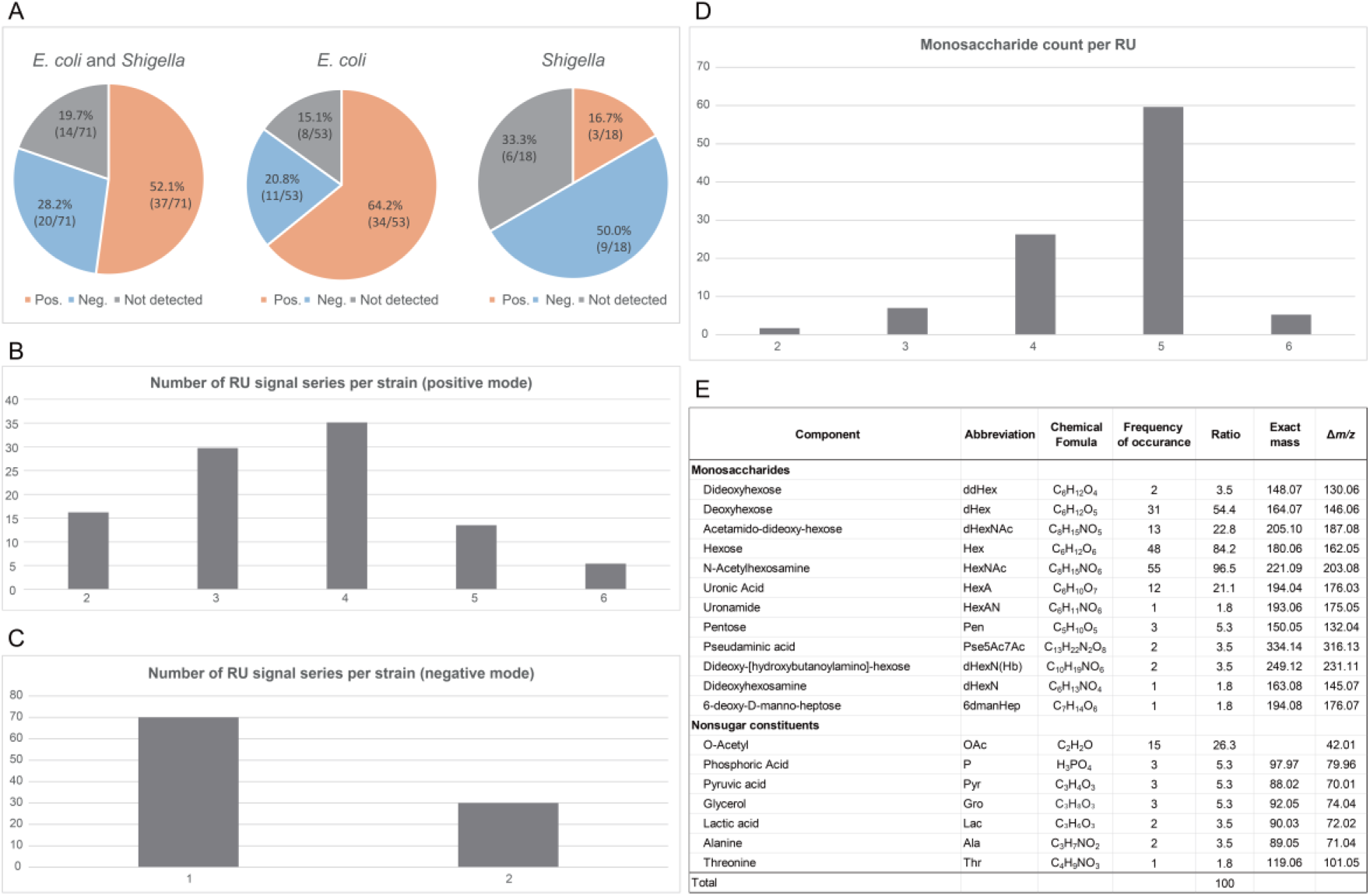
Species-scale validation of MALDI glycotyping across *Escherichia coli* and *Shigella* strains. (A) Detection of O-antigen repeating unit (RU)-derived signals across 71 strains analyzed by MALDI glycotyping. Pie charts indicate the proportion of strains yielding RU-derived signals in positive-ion mode (Pos), negative-ion mode (Neg), or lacking detectable signals (Not detected). Overall, RU signals were detected in 57 of 71 strains (>80%). (B and C) Number of RU-derived ion series detected per strain in positive-ion mode (B) and negative-ion mode (C). Most strains yielded multiple RU-derived ion series in positive-ion mode and one or two series in negative-ion mode. (D) Distribution of RU sizes inferred from MALDI-derived compositions, showing structures ranging from di- to hexasaccharides, with pentasaccharide RUs most frequently observed. (E) Monosaccharide and non-sugar constituents detected across all strains, together with their occurrence frequencies and exact masses, summarizing the compositional diversity of O-antigen repeating units identified by MALDI glycotyping.

These findings indicate that MALDI glycotyping enables scalable phenotypic characterization of O-antigen RU composition across diverse strains.

### MALDI glycotyping resolves modification-level structural variation among related O-antigens

To evaluate whether MALDI glycotyping can resolve modification-level variation among related O-antigens, we compared RU-derived spectral patterns across representative strain sets. Three independent O157 strains (GTC 03907, GTC 14600, and GTC 14561) yielded highly reproducible and nearly identical spectral patterns, demonstrating high spectral reproducibility for identical O-antigen phenotypes (<u>Fig. S3</u>).

In contrast, MALDI glycotyping readily resolved O-antigens with similar monosaccharide backbones but differing chemical modifications. For example, although GTC 03610 (O71 (20)) shares a closely related RU backbone with O157, distinct spectral patterns arose from differences in O-acetylation and stereochemistry, producing reproducible modification-dependent signal differences (Fig. 3A).

**Figure 3.**
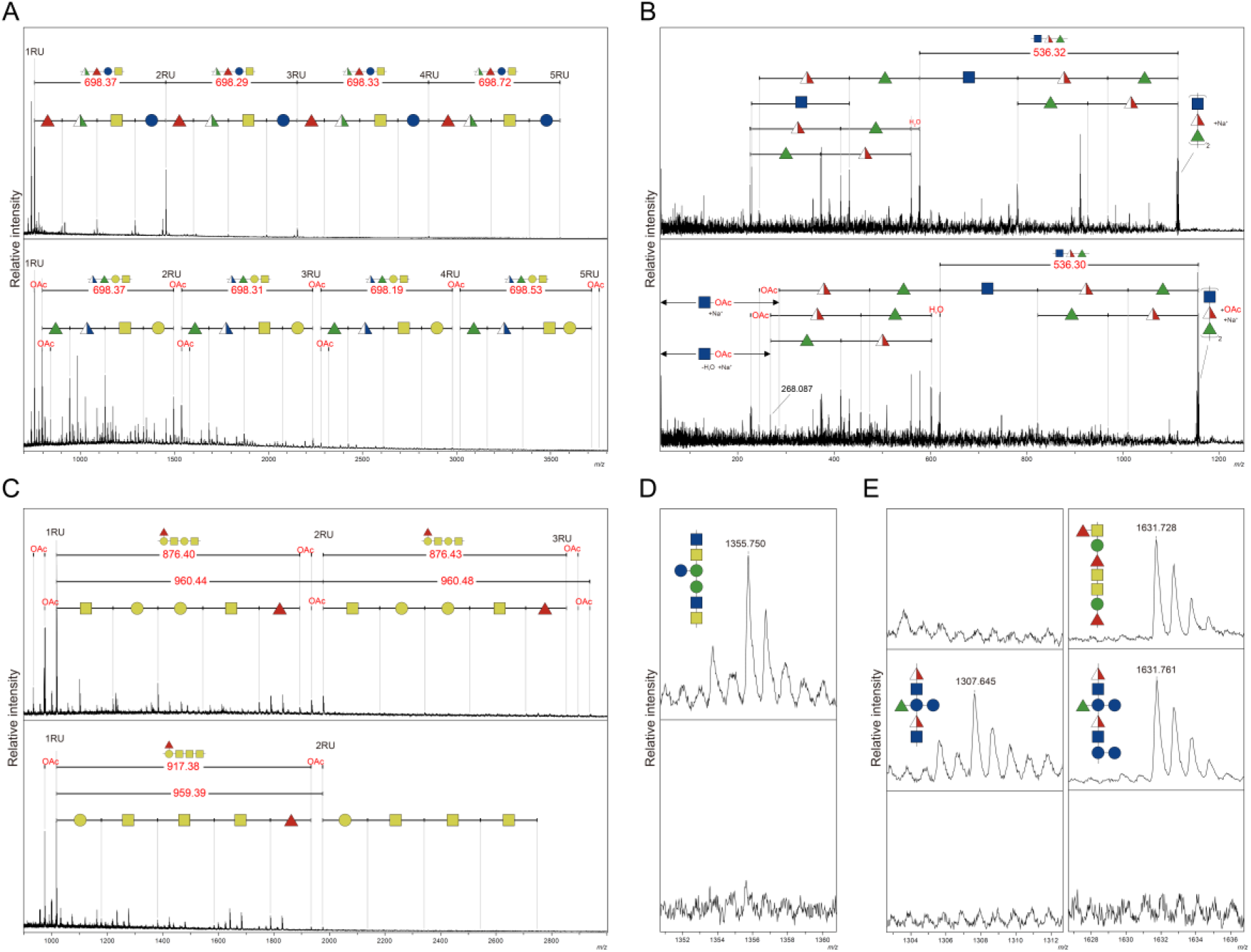
MALDI glycotyping resolves modification-level variation and discriminates isobaric O-antigen phenotypes. (A) Comparison of RU-derived MALDI spectra from *E. coli* GTC 03907 (O157, upper) and GTC 03610 (O71, lower). Although these strains share related repeating-unit (RU) glycan backbones, distinct spectral patterns are observed due to differences in O-acetylation and structural context. (B) Localization of O-acetylation sites by MS/MS analysis. MS/MS spectra of non-acetylated (*m/z* 1113.6) and monoacetylated (*m/z* 1155.6) 2RU precursor ions from *E. coli* GTC 14605 (O26) indicate O-acetylation of HexNAc residues within the trisaccharide RU. (C) Structural variation within a single serotype. Spectra from *E. coli* O128 strains GTC 14291 (upper) and GTC 13255 (lower) show differences in RU composition and O-acetylation state. (D) Discrimination of isobaric RUs. Expanded spectra of *E. coli* GTC 13256 (O6, upper) and GTC 03570 (O21, lower) share identical RU masses but display distinct spectral features, including a characteristic signal detected only in O6. (E) Comparison of isobaric RU phenotypes across multiple strains. Spectra from *E. coli* GTC 03561 (O11, upper), GTC 13262 (O25, middle), and GTC 03594 (O51, lower) exhibit distinct spectral patterns despite shared RU masses, reflecting differences in RU structural organization. Monosaccharide symbols follow the Symbol Nomenclature for Glycans (SNFG).

Sensitivity to partial modification was further illustrated by GTC 14605 (O26 (21)), which exhibited incomplete O-acetylation within a trisaccharide RU (HexNAc, dHexNAc, and dHex) (<u>Fig. S1.14</u>). MS/MS analysis of non-acetylated (*m/z* 1113.6) and monoacetylated (*m/z* 1155.6) 2RU precursor ions localized O-acetylation to HexNAc residues, providing MS/MS-based localization of a modification site not previously assigned for this O-antigen (Fig. 3B).

Differences in both RU sugar composition and modification state were similarly resolved among O128 variants. GTC 14291 and GTC 13255 differed by Δ*m/z* 1 per RU, consistent with hydroxyl-to-N-acetyl replacement and differences in O-acetylation state (Fig. 3C). MS/MS analysis further indicated colocalization of two O-acetyl groups on a hexose residue in GTC 14291, whereas the single O-acetyl group in GTC 13255 was assigned to either a hexose or HexNAc residue (<u>Fig. S4</u>).

Collectively, these results demonstrate that MALDI glycotyping provides reproducible, modification-sensitive spectral signatures that discriminate closely related O-antigens at the level of chemical modification and fine structural variation.

### Structure-dependent spectral signatures discriminate isobaric O-antigen repeating units

To evaluate whether RU-level phenotyping can discriminate O-antigens that are indistinguishable by exact mass alone, we analyzed isobaric RU pairs across the dataset (<u>Table S2</u>). Despite sharing identical RU masses, several pairs exhibited distinct MALDI spectral features, indicating structure-dependent spectral behavior beyond exact mass (<u>Fig. S5</u>).

For example, GTC 13256 (O6 (22)) and GTC 03570 (O21 (23)) both exhibited an RU mass series at Δ*m/z* 892 intervals yet produced distinct RU-derived spectral patterns. A characteristic signal associated with adjacent HexNAc residues was detected in O6 but absent in O21, providing a discriminating spectral feature (Fig. 3D<u>; Fig. S6A showing the reported O-antigen structures</u>). MS/MS analysis supported identical monosaccharide compositions, indicating that the observed differences arise from structural context rather than composition alone (<u>Fig. S7A</u>).

Similarly, strains GTC 03561 (O11 (24)), GTC 13262 (O25 (25)), and GTC 03594 (O51 (26)) shared an RU mass series at Δ*m/z* 860 intervals but displayed distinct MALDI spectral patterns. These spectral features reflected differences in compositional context and positional distribution of deoxyhexose residues within the O-antigen (<u>Fig. S6B showing the reported O-antigen structures</u>). Notably, a strain-specific deoxyHexNAc-associated signal was uniquely observed in O25, consistent with its reported RU composition (Fig. 3E). In addition, deoxyhexoses located in side chains in O11 and O25 were susceptible to acid treatment, whereas deoxyhexoses positioned in the main chain in O51 remained intact, accounting for the observed spectral divergence. MS/MS analysis provided representative structural support for these interpretations (<u>Fig. S7B</u>).

These results demonstrate that MALDI glycotyping can discriminate isobaric O-antigen RUs by capturing structure-dependent spectral features beyond exact mass.

### Resolution of ambiguous and cross-reactive O-antigen phenotypes

To examine how MALDI glycotyping performs in challenging analytical situations encountered during O-antigen characterization, we analyzed representative cases of ambiguous spectral patterns, cross-reactive antigens, and intra-serotype variants.

In some strains, positive-ion mode spectra produced repeating hexose-derived ion series that were insufficient for assignment based solely on RU composition (<u>Fig. S8</u>). For representative antigens such as O62 (27), characteristic RU spectral patterns supported the reported pentasaccharide hexose structure. In contrast, several non-hexose O-antigens required sequential negative-ion mode acquisition to detect compositionally informative RU-derived signals (ex. GTC 03612, GTC 03638, and GTC 13248; <u>Fig. S1.33, S1.36, and S1.44</u>). This conditional dual-ion mode workflow enabled detection of RU-derived signals that were not detectable in positive-ion mode alone, thereby resolving otherwise ambiguous spectral patterns.

To assess how MALDI glycotyping behaves in cross-reactive contexts, we next examined reported *Shigella*–*Escherichia coli* O-antigen pairs that share antigenic determinants (17). These pairs yielded highly similar MALDI spectral patterns consistent with shared RU architectures across species and their known serological cross-reactivity (<u>Fig. S9 and Table S3</u>). Nevertheless, modification-level differences, such as O-acetylation, remained detectable in the MALDI spectra.

We also evaluated strains for which no O-antigen–derived signals were detected by MALDI glycotyping (14 strains in total). Complementary antiserum testing was performed for 9 of these strains. The remaining five strains were not tested due to lack of corresponding antisera availability. Two *E. coli* strains (GTC 13261 and GTC 13305) were negative by antisera despite their registered O-antigen assignments, indicating that the corresponding O-antigens were not expressed under the tested conditions rather than representing analytical failure (<u>Fig. S10</u>).

Finally, intra-serotype variants were examined across eight O-antigen groups by comparing reported RU compositions with those inferred by MALDI glycotyping (<u>Table S4</u>). MALDI glycotyping enabled discrimination of multiple variants within these serotypes, with MS/MS analysis providing structural support for representative strains (<u>Fig. S11</u>).

Collectively, these results show that MALDI glycotyping resolves analytical ambiguity in RU detection, reflects known cross-reactive O-antigen relationships across species, and enables discrimination of intra-serotype structural variants at the RU level.

### Resolution of discrepant O-antigen assignments by integrated MALDI glycotyping and serology

To investigate discrepancies between MALDI glycotyping phenotypes and registered O-antigen assignments, we analyzed eight strains whose MALDI glycotyping spectra were inconsistent with their reported O-antigen labels. MS and MS/MS analyses revealed RU structural features and sugar compositions inconsistent with previously reported O-antigen references (<u>Fig. S12.1–S12.8 and Table S5</u>).

To exclude capsular (K) antigens as potential sources of the detected RU signals, the observed RU masses were compared with reported K-antigen structures (18). None of the strains exhibited RU compositions consistent with known K-antigens (<u>Table S6</u>).

Agglutination serotyping was performed using a commercial antisera panel covering 50 *E. coli* O-antigen types. All strains for which corresponding antisera were available were negative in agglutination tests despite their registered O-antigen assignments (<u>Fig. S13</u>).

Subsequent targeted serological screening using the antisera panel identified alternative O-antigen types for four strains (<u>Fig. S14</u>). These assignments were fully consistent with RU compositions and spectral features determined by MALDI glycotyping (Fig. 4<u>; Table S7</u>). These findings demonstrate that MALDI glycotyping can identify potential inconsistencies in archived serotype annotations and guide targeted confirmatory serological testing.

**Figure 4.**
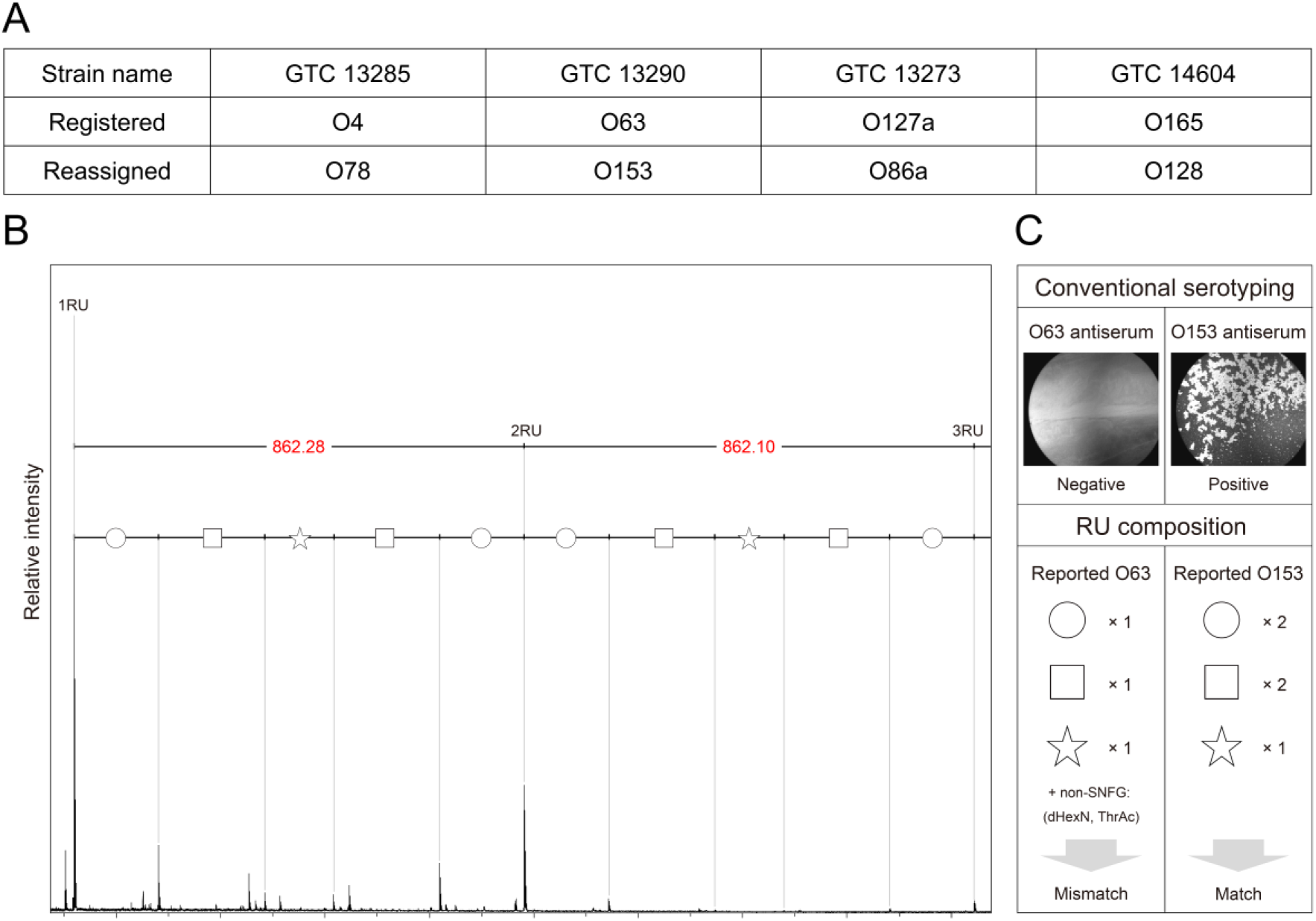
Resolution of discrepant O-antigen assignments by integrated MALDI glycotyping and serology. (A) Summary of four *E. coli* strains (GTC 13285, GTC 13290, GTC 13273, and GTC 14604) in which registered O-antigen assignments were discordant with MALDI glycotyping phenotypes. Agglutination testing reassigned these strains to O78, O153, O86a, and O128, respectively. (B) Representative example of discordance between registered serotype and MALDI glycotyping phenotype. The MALDI spectrum of *E. coli* GTC 13290 shows RU-derived ion series inconsistent with the reported RU composition of O63. The observed RU mass (Δ*m/z* ≈ 862) and inferred composition (Hex₂HexNAc₂Pent₁) are consistent with the reported O153 O-antigen structure. (C) Serological confirmation of MALDI-based reassignment for strain GTC 13290. Agglutination testing shows no reactivity with O63 antiserum and clear reactivity with O153 antiserum. A schematic comparison of MALDI-derived RU composition with reported O-antigen structures highlights mismatch with O63 and agreement with O153.

Independent comparison of MALDI-derived RU compositions and exact masses with reported *E. coli* O-antigen structures (<u>Table S8.1–S8.4</u>) further constrained candidate identities for the reassigned strains. Two strains uniquely matched alternative O-antigen types, whereas two corresponded to multiple isobaric candidates <u>(Table S9A</u>). In all cases, MALDI-based structural inference was concordant with agglutination results.

Analysis of reassigned O128 strains further showed that differences in O-acetylation state corresponded to differences in serological reactivity, linking RU structural variation with antigenic reactivity (<u>Fig. S15 and Table S10</u>).

The remaining strains that showed no agglutination with the antisera panel were therefore evaluated based solely on MALDI glycotyping–derived RU compositions and spectral features to infer compatible O-antigen candidates.

Together, these results demonstrate that discordance between archived serotype labels and expressed O-antigen phenotypes can occur even in curated strain collections, and that integration of MALDI glycotyping with serology enables resolution of such discrepancies.

### Inference of unresolved O-antigen identities for serologically untypeable strains

To examine how MALDI glycotyping performs in cases unresolved by serology, we analyzed four *E. coli* strains (GTC 13263, GTC 13282, GTC 13293, and GTC 13264) that showed no detectable agglutination with a commercial antisera panel despite having reported O-antigen assignments (<u>Fig. S14</u>).

Candidate O-antigen identities were first constrained by agreement between RU exact masses and reported *E. coli* O-antigen structures (Fig. 5). Candidate sets were then further evaluated by comparison of MALDI-derived RU sugar compositions and observed modification states (<u>Table S9B</u>).

**Figure 5.**
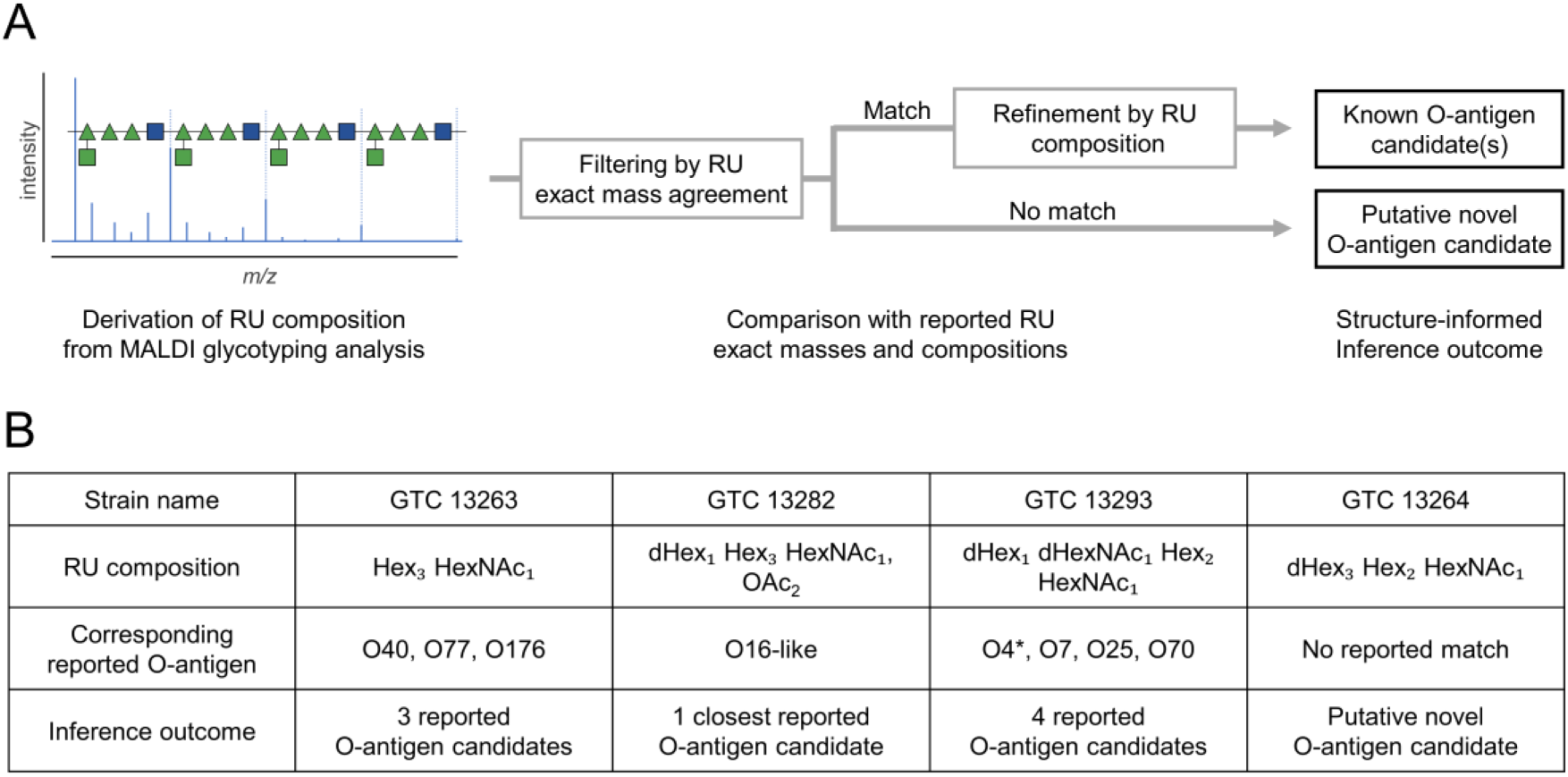
Inference of unresolved O-antigen identities using MALDI glycotyping. (A) Schematic overview of the inference workflow used to interpret strains unresolved by serological testing. MALDI glycotyping provides RU-derived phenotypic features, including exact mass and inferred sugar composition. Candidate O-antigen identities were first constrained by agreement between estimated and reported RU exact masses and subsequently refined using MALDI-derived RU composition and observable modification states. (B) Application of the inference framework to four *E. coli* strains that showed no agglutination with the antisera panel. MALDI-derived RU compositions restricted candidate identities to several reported O-antigens (GTC 13263 and GTC 13293), suggested a closest reported O-antigen match (GTC 13282), or showed no correspondence with reported structures (GTC 13264). O4* denotes Jansson’s O4 variant.

For GTC 13263, the observed RU mass matched reported O-antigen structures with the same sugar composition (three Hex and one HexNAc). These included several reported O-antigen types, such as O40, O77, and O176, preventing unique assignment but restricting the candidate identities to a small set of compatible structures (<u>Fig. S16A</u>).

Similarly, GTC 13282 exhibited an RU mass matching reported O-antigen structures with the same sugar composition excluding O-acetylation (one dHex, three Hex, and one HexNAc), which was consistent with both O16 and O18 (<u>Fig. S16B</u>). However, MALDI spectra indicated an O-acetylation state not matching reported RU structures. Because the reported O16 RU shares the same sugar composition with a single O-acetylation, this strain was interpreted as an O16-like structural variant with an altered O-acetylation state. Lack of reactivity to anti-O18 serum further supported this interpretation.

For GTC 13293, the observed RU mass was shared by multiple reported O-antigens (<u>Fig. S16C</u>). Comparison of MALDI-derived RU sugar composition further restricted the candidates to four reported O-antigen structures, such as Jansson’s O4, O7, O25, and O70. Analysis of characteristic MALDI spectral patterns, including the presence of a [2RU − Hex + K]^+^ signal and absence of the corresponding [2RU − dHex + K]^+^ signal (<u>Fig. S17</u>), suggested hexose-only side chains and further restricted compatible structures to Jansson’s O4 or O70.

In contrast, the RU composition observed for GTC 13264 (three dHex, two Hex, and one HexNAc) did not match any previously reported *E. coli* O-antigen RU composition, suggesting the presence of either an uncharacterized structural variant or a previously unreported O-antigen phenotype.

These results demonstrate that MALDI glycotyping enables RU-level structural inference that can restrict candidate O-antigen identities and identify potentially uncharacterized O-antigen phenotypes when serological typing is inconclusive.

## Discussion

Rapid and reliable O-antigen characterization remains important for clinical microbiology, where serotype information supports pathogen surveillance, outbreak investigations, and quality assurance of reference strain collections. In this study, we establish MALDI glycotyping as a rapid phenotypic readout of O-antigen repeating units (RUs) in *Escherichia coli* and *Shigella*, extending routine MALDI-TOF MS workflows beyond protein fingerprinting to enable surface glycan phenotyping.

A key feature of the workflow is sequential dual-ion mode acquisition, in which negative-ion mode is triggered only when RU signals are absent in positive-ion mode. This conditional acquisition strategy enables detection of acidic O-antigen phenotypes while minimizing ion-mode-dependent detection bias. Applied to a diverse panel of 71 *E. coli* and *Shigella* strains, RU-derived signals were detected in more than 80% of strains, supporting broad phenotypic coverage under routine laboratory conditions. The remaining strains lacking detectable RU signals likely reflect several factors, including biological variability such as rough phenotypes lacking O-antigens or low expression levels, as well as O-antigen structures that exhibit lower ionization efficiency under the current experimental conditions. Notably, several strains without detectable RU signals were also negative in agglutination despite their registered serotype annotations, supporting biological absence or low expression of O-antigens rather than analytical failure of the MALDI glycotyping workflow. Despite these cases of non-detection, the detected RU phenotypes captured substantial compositional diversity and produced interpretable spectral signatures reflecting modification-level variation and structural context. These features enabled discrimination of closely related O-antigens, modification-level variants, and isobaric RUs. This study therefore substantially expands the scale of evaluation compared with earlier implementations, which were tested on only a limited number of strains.

From a clinical perspective, one of the most important findings is that discordance between archived serotype labels and expressed O-antigen phenotypes can occur even in curated strain collections. Eight strains showed MALDI phenotypes inconsistent with their registered O-antigen assignments, and integrated agglutination testing enabled reassignment of four strains to alternative O-antigen types fully consistent with MALDI-derived RU compositions. These observations support MALDI glycotyping as a practical quality-control layer that can flag potential inconsistencies for targeted follow-up and help contextualize archived serotype annotations with direct phenotypic evidence. Such phenotype-based confirmation may be particularly valuable in epidemiological investigations where accurate serotype annotation is critical for strain tracking and outbreak analysis (28). More broadly, phenotype-based O-antigen confirmation may help improve the reliability of archived strain annotations used in reference collections and surveillance datasets.

Similarly, some strains without detectable RU signals also lacked agglutination despite their registered O-antigen assignments. This observation suggests that factors such as strain provenance, biological variability, or handling history may contribute to discordance between archived serotype labels and expressed O-antigen phenotypes. Thus, non-detection is often biologically interpretable and can itself provide useful information for follow-up testing.

Notably, MALDI glycotyping provides RU-level phenotypes rather than forced serotype calls (12, 29). We therefore implemented a structure-informed interpretation strategy in which RU exact mass, inferred composition, and modification states constrain candidate O-antigen identities for strains unresolved by serology. This framework effectively narrows candidate space and highlights cases lacking correspondence with reported structures that may represent uncharacterized glycotypes. Comparison with a compiled reference landscape of reported O-antigen RU compositions and exact masses further supports systematic contextualization of MALDI-derived phenotypes within known structural repertoires.

Comprehensive serological coverage of *E. coli* O-antigens is intrinsically limited in routine laboratories. Although more than 180 O-antigen types have been described, commercially available antisera panels typically cover only a subset of these serotypes, making complete serological characterization difficult in practice. In contrast, MALDI glycotyping detects O-antigen structural signatures directly and therefore does not rely on predefined antisera panels. This feature enables broader and potentially more comprehensive phenotypic characterization of O-antigen diversity while remaining compatible with routine diagnostic workflows. Although multiple colonies were used here to standardize cell density, the workflow remains compatible with single-colony sampling similar to that used in routine MALDI identification workflows (12). In principle, species identification by protein fingerprinting and O-antigen phenotyping by MALDI glycotyping could be performed using the same instrument and target plate with differences only in sample pretreatment.

This compatibility may facilitate practical adoption of O-antigen phenotyping in laboratories already equipped with MALDI-based identification systems. Importantly, MALDI glycotyping provides direct structural information on expressed O-antigens, offering a phenotypic dimension beyond that provided by serological typing. As MALDI-TOF MS instruments are already widely implemented in clinical microbiology laboratories, extension of these platforms to O-antigen phenotyping may provide a practical route for expanding phenotypic characterization without requiring major infrastructure changes (30, 31).

This study has limitations. RU detection remains dependent on the specific chemotype and the unified pretreatment parameters used. Further optimization of hydrolysis conditions may improve coverage for particularly recalcitrant structures. Additionally, our serological follow-up was limited by antiserum availability, and the accuracy of structural inference remains dependent on the completeness of curated reference databases. Future expansion of spectral libraries will be essential to reduce ambiguity in compositionally similar contexts.

In conclusion, MALDI glycotyping provides a robust, rapid, and infrastructure-compatible approach for O-antigen characterization. By integrating this method with targeted serology, clinical laboratories can resolve discrepancies in strain identification and gain a deeper understanding of expressed bacterial diversity. Because the workflow requires only brief acid pretreatment and utilizes the same MALDI-TOF MS platforms already ubiquitous in clinical settings, it offers a practical and high-resolution complement to traditional typing methods, facilitating the integration of glycan phenotyping into routine clinical microbiology.

## ACKNOWLEDGMENTS

This research is supported by A-STEP (JPMJTM20JB to H.H.) from the Japan Science and Technology Agency (JST), Grant-in-Aid for Scientific Research (B: 22H02191, 23K23458 to H.H.), Core-to-Core type B (JPJSCCB20240004 to H.H.), Grant-in-Aid for JSPS Fellows (23KJ0052 to S.U.) from the Japan Society for the Promotion of Science (JSPS). We thank the Gifu Type Culture Collection of the Microbial Genetic Resource Stock Center, Gifu University Graduate School of Medicine, for providing bacterial strains through the National BioResource Project (NBRP), Japan.

## Contributions

H.H. conceived and supervised the project; S.U. performed the experiments; S.U. and H.H. analyzed the data and wrote the paper.

